# EsaB is a core component of the *Staphylococcus aureus* Type VII secretion system

**DOI:** 10.1101/151316

**Authors:** M. Guillermina Casabona, Grant Buchanan, Martin Zoltner, Catriona P. Harkins, Matthew T.G. Holden, Tracy Palmer

**Affiliations:** Division of Molecular Microbiology School of Life Sciences, University of Dundee, Dundee, UK.; School of Medicine, University of St Andrews, St Andrews, KY16 9TF, UK.

**Keywords:** *Staphylococcus aureus*, Protein secretion, T7SS, Regulation

## Abstract

Type VII secretion systems (T7SS) are found in many bacteria and secrete proteins involved in virulence and bacterial competition. In *Staphylococcus aureus* the small ubiquitin-like EsaB protein has been previously implicated as having a regulatory role in the production of the EsxC substrate. Here we show that in the *S. aureus* RN6390 strain, EsaB does not genetically regulate production of any T7 substrates or components, but is indispensable for secretion activity. Consistent with EsaB being a core component of the T7SS, loss of either EsaB or EssC are associated with upregulation of a common set of iron acquisition genes. However, a further subset of genes were dysregulated only in the absence of EsaB. In addition, fractionation revealed that although an EsaB fusion to yellow fluorescent protein partially localised to the membrane, it was still membrane-localised when the T7SS was absent. Taken together our findings suggest that EsaB has T7SS-dependent and T7SS-independent roles in *S. aureus*.

## INTRODUCTION

Protein secretion systems are nanomachines employed by bacteria to transport protein substrates across their cell envelopes. Gram-negative bacteria produce a number of different secretion machineries that export proteins involved in a wide variety of processes including signalling, nutrient scavenging, host interaction and virulence (1). The type VII secretion system (T7SS) is found in some Gram-negative and many Gram-positive bacteria, and is particularly common among organisms of the actinobacteria and firmicutes phyla (2). The T7SS was initially described in the pathogenic mycobacteria *Mycobacterium tuberculosis* and *Mycobacterium bovis,* where the ESX-1 T7SS was shown to be essential for virulence, due to the secretion of two major T-cell antigens EsxA (formerly known as ESAT-6) and EsxB (formerly known as CFP-10) (3-5). EsxA and EsxB are founding members of the WXG100 protein family that appear to be exclusively linked to T7SSs, and all characterised T7 systems are associated with at least one family member. The presence of a membrane-bound ATPase of the SpoIIIE/FtsK family (termed EccC in actinobacteria and EssC in firmicutes) is another hallmark of all T7SSs (6). In Mycobacteria, three further membrane proteins EccB, EccD and EccE assemble with EccC to form a large 1.5 MDa core complex (7, 8). This complex further associates with a membrane-bound mycosin serine protease, MycP, that is essential for T7 protein secretion and for stability of the membrane complex (9).

*Staphylococcus aureus*, an opportunistic pathogen of humans and animals, also elaborates a T7SS that is distantly related to the T7SSs found in mycobacteria (10). Mutational analysis has indicated that it plays an important role in persistence in mouse models of infection, intraspecies competition and potentially iron homeostasis (10-15). In commonly-studied strains of *S. aureus* such as Newman, USA300 and RN6390, the secretion system is encoded by the 12 gene *ess* locus (10, 12, 16). The first six genes at this locus encode core components of the secretion machinery, including the WXG100 protein EsxA and the SpoIIIE/FtsK ATPase EssC (Fig 1A,B). However, *S. aureus* and other firmicutes lack homologues of EccB, EccD, EccE and MycP and instead have an apparently unrelated set of membrane-bound secretion components (EsaA, EssA and EssB in *S. aureus*) (12, 17-19). The sixth component of the *S. aureus* T7SS is EsaB, which is predicted to be a small cytoplasmic protein of 80 amino acids that is structurally related to ubiquitin (20). In *S. aureus* strains Newman and USA300, a transposon insertion in *esaB* does not abolish secretion of T7 substrates but is linked with an increase in RNA transcripts covering the gene encoding the substrate EsxC (11). By contrast, in-frame deletion of *esaB* abolished EsxA and EsxC secretion in strain RN6390 but did not detectably affect production of these substrate proteins (12). Similarly, inactivation of *yukD*, which encodes the *Bacillus subtilis esaB* homologue, also abolished T7 secretion (17, 18).

**Figure 1.**
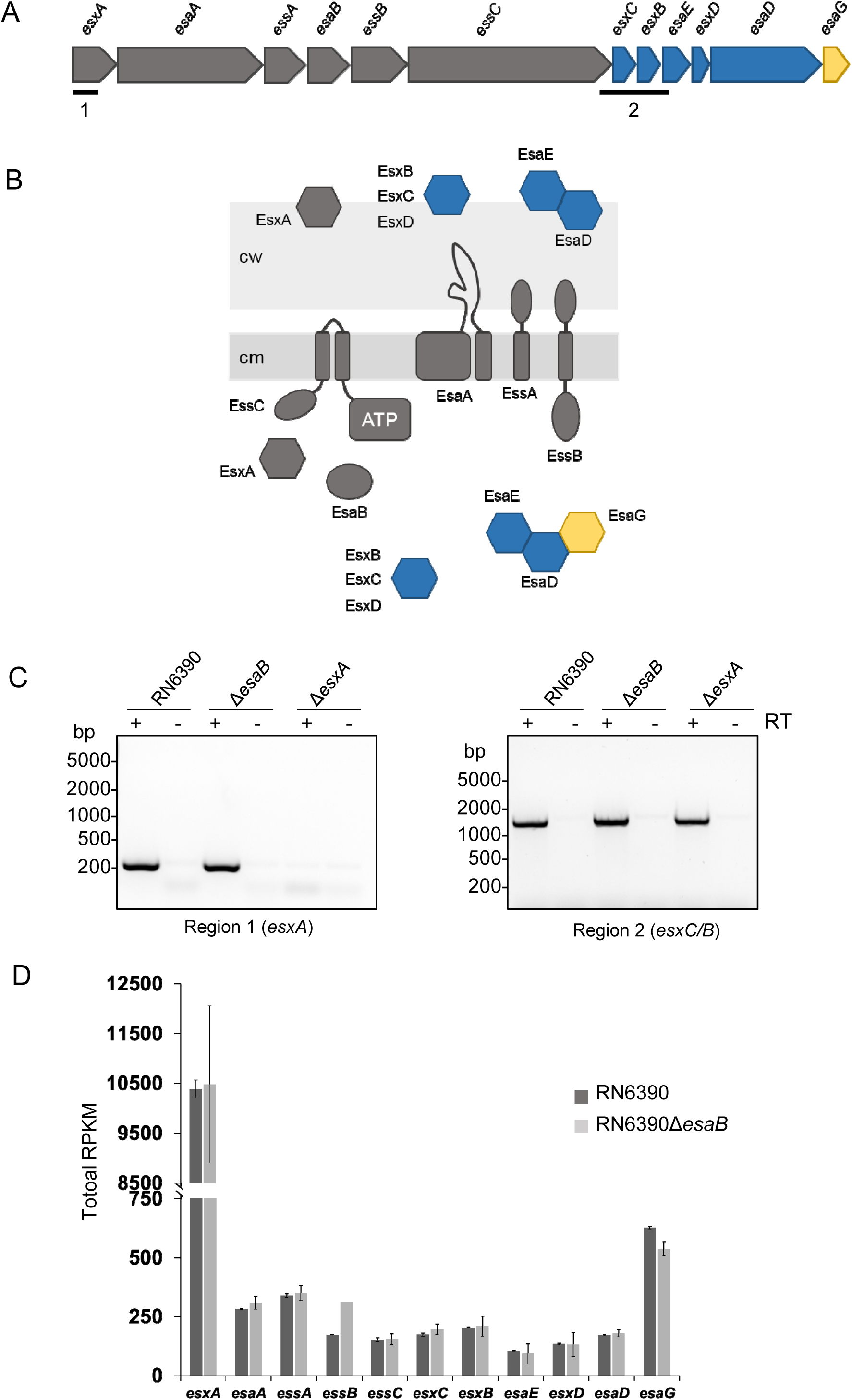
EsaB is not a transcriptional regulator. (A) The *ess* locus in *S. aureus* RN6390. Genes encoding core components are shaded in grey, secreted proteins in blue and a cytoplasmic antitoxin in yellow. The regions analysed by RT-PCR are indicated. (B) Predicted subcellular locations of Ess-encoded components. cw – cell wall, cm – cytoplasmic membrane. (C) RT-PCR analysis of *esxA* (region 1) and *esxC/B* (region 2) from the RN6390 and isogenic *esaB* and *esxA* mutant strains. Equivalent amounts of mRNA from each strain were used to generate cDNA. RT: reverse transcriptase. (D) Total mRNA counts of *ess* genes from RNA-Seq analysis of RN6390 and the *esaB* mutant strain. RPKM - reads of transcript per kilobase per million of mapped reads.

In this study, we have addressed the role of EsaB in *S. aureus* T7 secretion using strain RN6390. We show that EsaB does not regulate *esxC* transcripts or those of other *ess*-encoded genes. Instead our findings show that EsaB behaves as a core component of the T7SS. Interestingly, however, RNA-Seq analysis identified a subset of genes from the AirSR regulon that showing altered regulation in the absence of EsaB, suggesting that it may play additional, T7SS-independent roles in *S. aureus* physiology.

## METHODS

### Bacterial strains and growth conditions

*S. aureus* strain RN6390 (NCTC8325 derivative, *rbsU*, *tcaR*, cured of (ϕ11, (ϕ12, (ϕ13; (21)) and the isogenic Δ*esaB* and Δ*esx* (Δ*esxA* – *esaG*) strains (12) were employed in this study. *S. aureus* strains were cultured in Tryptic Soy Broth (TSB) at 37°C with shaking unless otherwise stated. For calculation of cell numbers we estimated by dilution analysis that one unit at OD 600nm corresponds to 6x10^8^ CFU for strain. When required, chloramphenicol (Cml, final concentration 10 μg/ml) was added for plasmid selection. *E. coli* strain JM110 (Stratagene) was used for cloning purposes and BL21(DE3) (22) for EsaB overproduction and purification. *E. coli* was grown in Luria-Bertani (LB) medium at 37°C with agitation. When appropriate, ampicillin was used for plasmid selection (final concentration 125 μg/ml).

### Genetic constructs

All plasmids used in this study are listed in Table 1. The *esaB* gene with its own RBS was PCR amplified from *S. aureus* RN6390 genomic DNA using primers EsaB-fw and EsaB-rev (Table S1). The 0.3 kb *Hpa*I/*Eco*RI restriction fragment was cloned into pRAB11 under control of the tetracycline inducible promoter, giving pRAB11-esaB. Clones were selected in *E. coli* and verified by DNA sequencing. Plasmid pRAB11-esaB-YFP was generated by cloning the 0.3 kb *Hpa*I/*Eco*RI restriction fragment into pRAB11-YFP (15). Clones were selected in *E. coli* and verified by DNA sequencing. Nucleotide variants of *esaB* were generated by the Quickchange site-directed mutagenesis protocol (Stratagene) using pRAB11-esaB or pRAB11-esaB-YFP as a template and primers listed in Table S1. Modified plasmids were digested using *Dpn*I for at least 1h at 37°C and transformed into *E. coli*. Single point mutations were verified by DNA sequencing.

**Table 1.** Plasmids used in this study.

| Plasmid | Relevant genotype or description | Source or reference |
| --- | --- | --- |
| pRAB11 | *E. coli*/*S. aureus* shuttle vector, inducible protein expression, Amp<sup>r</sup>, Cml<sup>r</sup> | (35) |
| pRAB11-esaB | pRAB11 producing EsaB | This study |
| pRAB11-esaB-V7A | pRAB11 producing V7A-substituted EsaB | This study |
| pRAB11-esaB-V7K | pRAB11 producing V7K -substituted EsaB | This study |
| pRAB11-esaB-T8A | pRAB11 producing T8A-substituted EsaB | This study |
| pRAB11-esaB-T8E | pRAB11 producing T8E-substituted EsaB | This study |
| pRAB11-esaB-T8R | pRAB11 producing T8R-substituted EsaB | This study |
| pRAB11-esaB-T8H | pRAB11 producing T8H-substituted EsaB | This study |
| pRAB11-esaB-T8K | pRAB11 producing T8K-substituted EsaB | This study |
| pRAB11-esaB-T8S | pRAB11 producing T8S -substituted EsaB | This study |
| pRAB11-esaB-D10A | pRAB11 producing D10A-substituted EsaB | This study |
| pRAB11-esaB-D20A | pRAB11 producing D20A-substituted EsaB | This study |
| pRAB11-esaB-L21A | pRAB11 producing L21A-substituted EsaB | This study |
| pRAB11-esaB-K30A | pRAB11 producing K30A-substituted EsaB | This study |
| pRAB11-esaB-I44A | pRAB11 producing I44A-substituted EsaB | This study |
| pRAB11-esaB-I44K | pRAB11 producing I44K-substituted EsaB | This study |
| pRAB11-esaB-K52A | pRAB11 producing K52A-substituted EsaB | This study |
| pRAB11-esaB-K56A | pRAB11 producing K56A-substituted EsaB | This study |
| pRAB11-esaB-L66A | pRAB11 producing L66A-substituted EsaB | This study |
| pRAB11-esaB-I71A | pRAB11 producing I71A-substituted EsaB | This study |
| pRAB11-esaB-I71K | pRAB11 producing I71K -substituted EsaB | This study |
| pRAB11-esaB-G74A | pRAB11 producing G74A-substituted EsaB | This study |
| pRAB11-esaB-D75A | pRAB11 producing D75A-substituted EsaB | This study |
| pRAB11-esaB-L77A | pRAB11 producing L77A-substituted EsaB | This study |
| pRAB11-esaB-L77K | pRAB11 producing L77K -substituted EsaB | This study |
| pRAB11-esaB-YFP | pRAB11 producing EsaB-YFP | This study |
| pRAB11-esaB-V7K-YFP | pRAB11 producing V7K-substituted EsaB-YFP | This study |
| pRAB11-esaB-T8A-YFP | pRAB11 producing T8A-substituted EsaB-YFP | This study |
| pRAB11-esaB-T8R-YFP | pRAB11 producing T8R-substituted EsaB-YFP | This study |
| pRAB11-esaB-I71K-YFP | pRAB11 producing I71K-substituted EsaB-YFP | This study |
| pET15b-HISEsaB | pET15b expressing 6XHis-tagged EsaB | This study |

### RNA isolation and RT-PCR

For RNA-Seq analysis, three biological repeats of the *S. aureus esaB* strain was grown aerobically in TSB up to an OD_600_ of 1 at which point mRNA was prepared (in three technical replicates). This experiment was carried out alongside the RN6390 and *essC* strains (15) and followed identical methodology.

For RT-PCR, the indicated *S. aureus* strains were grown aerobically in TSB and harvested at an OD_600_ of 1. At this point, the mRNA was extracted using the SV total RNA Isolation Kit (Promega) with some minor modifications. Cell samples were stabilized in 5% phenol/95% ethanol on ice for at least 30 min and then centrifuged at 2770 *g* from 10 min. Cells were then resuspended in 100 μl of TE buffer containing 500 μg ml^-1^ lysostaphin and 50 μg ml^-1^ lysozyme and incubated at 37°C from 30 min. Subsequently, the manufacturer’s instructions were followed. Isolated RNA was subjected to a second DNase treatment using the DNA-free kit (Ambion). RNA was stored at -80°C until use. RT-PCR to probe transcription of genes in the indicated strains was carried out using 500 ng of mRNA as template with the indicated primers (Table S1). PCR products were visualized on 1% agarose gels.

### Purification of 6His-EsaB and generation of polyclonal antisera

The EsaB coding sequence (UniProt code ESAB_STAAM) was PCR amplified from a synthetic gene (codon optimized for *Escherichia coli* K12 (Genscript)) using the primers EsaB-pET1 and EsaB-pET2 (Table S1) and cloned into the *Nde*I*/Xho*I site of a modified pET15b vector (Novagen). The plasmid produces an N-terminal His_6_-tagged protein with a TEV (tobacco etch virus) protease cleavage site. The protein was expressed and purified as described previously (23), except the tag-free EsaB was not collected in the flow-through of the negative purification but required a 30mM imidazole elution. The final size exclusion chromatography step used a 24ml HR 30/100 GL Superdex75 column (GE healthcare), equilibrated with 20 mM Tris pH 7.8, 100 mM NaCl and was calibrated with molecular mass standards (thyroglobulin, 670 kDa; γ-globulin, 158 kDa; serum albumin, 67 kDa; ovalbumin; 44 kDa, myoglobin, 17 kDa; and vitamin B12, 1 kDa). 2 mg purified EsaB (retaining a Gly–Ala–Ser–Thr sequence at the N-terminus after the cleavage step) was utilized as antigen to immunize rabbits for polyclonal antibody production in a standard three injections protocol (Seqlab, Goettingen, Germany).

### Secretion assays, subcellular fractionation and western blotting

The indicated strains were grown overnight in TSB, diluted 1/100 in fresh medium and grown up to mid-log phase, at which point whole cells and supernatant fractions were harvested as described previously (12). Briefly, cells and supernatant were separated by a 10 min centrifugation step at 2770 *g*. Cells were washed twice with PBS, adjusted to and OD_600_ of 1 and digested using 50 μg/ml of lysostaphin by incubation at 37°C for 30 min. Supernatants were filtered using a 0.22 μm filter and TCA-precipitated in the presence of 50 μg/ml deoxycholate, as described. For *S. aureus* subcellular fractionation, cells were grown to mid-log phase with shaking and treated as previously described (12). Briefly, cells were harvested by centrifugation and resuspended in TSM buffer (50 mM Tris-HCl pH 7.6, 0.5 M sucrose, 10 mM MgCl_2_). Lysostaphin was added to a final concentration of 50 μg ml^-1^ and cells were incubated at 37°C for 30 min to digest the cell wall. At this point, protoplasts were sedimented to recover the cell wall (supernatant fraction). Protoplasts were disrupted by sonication and the membrane was obtained after an ultracentrifugation step at 227 000 *g* for 30 min and at 4°C. The supernatant was retained as the cytoplasmic fraction. Samples were boiled for 10 min prior to separation in bis-Tris gels and subsequent western blotting.

Polyclonal antisera were used at the following dilutions: α-EsxA 1:2500 (12), α-EsxB 1:1000 (15), α-EsxC 1:2000 (12), α-EsaB 1:500, α-TrxA 1:20000 (24) and α-SrtA (Abcam) 1:3000. Anti-GFP antibody was obtained from Roche and used according to manufacturer’s instructions.

## RESULTS

### EsaB does not regulate the level of *esxC* transcripts in strain RN6390

A previous study has shown that a transposon insertion in the *esaB* gene results in an increase in *esxC* transcripts in the Newman and USA300 strain backgrounds, and a concomitant increase in the EsxC polypeptide, implicating it as a regulator (11). To investigate whether loss of *esaB* by in-frame deletion affects the level of *esxC* mRNA in strain RN6390, we isolated mRNA from the parental strain and the isogenic *esaB* mutant, prepared cDNA and undertook reverse transcriptase PCR with primers covering either *esxA* (the first gene at the *ess* locus, included as a negative control) or *esxC* (Fig 1A). It can be seen (Fig 1C) that the level of transcripts for each of these genes was qualitatively similar in the wild type and *esaB* backgrounds.

To examine this quantitatively, we undertook RNA-Seq analysis on RNA prepared from three biological repeats of the RN6390 and *esaB* strains grown aerobically in TSB to an OD_600_ of 1. Note that these experiments were performed at the same time as the RN6390 vs *essC* RNA-Seq analysis described in (15) and used the same RN6390 dataset. Fig 1D shows that the level of *esxC* transcripts were indistinguishable between the wild type and *esaB* strains. Analysis of the transcript levels of the other genes at the *ess* locus indicates that in general they were also not significantly altered by the loss of *esaB* although there was a small increase in the level of *essB*. We conclude that there is no evidence that *esaB* regulates the level of *esxC* transcripts in RN6390.

We next examined the entire transcript profile of the *esaB* mutant to investigate the transcriptional/post-transcriptional response to the loss of this small protein. We found 101 genes de-regulated in the *esaB* mutant compared to the parental strain (using a cut off of logFC > 2 or < -2 and qvalue < 0.05, as applied previously (15)), Fig 2A. Of these, 43 were upregulated by the loss of *esaB* whereas 58 were downregulated when *esaB* was absent – these genes are listed in Table 2. Interestingly, almost all of the genes that were differentially regulated in the *essC* mutant (15) were also similarly regulated in the *esaB* strain (Fig 4B), although there was a substantive subset of genes that were differentially expressed in the *esaB* mutant but not the *essC* strain (Table 2). It can be seen that almost all of the iron acquisition genes, including those for heme acquisition, staphyloferrin synthesis and uptake and ferrichrome import were commonly upregulated by loss of either *esaB* or *essC* (Table 2). Furthermore six of the eight downregulated genes from the *essC* strain were also down regulated in the *esaB* strain (note that one of the two genes unaffected in the *esaB* dataset is *essC* itself, which appears downregulated in the *essC* dataset because it has been deleted). The finding that almost the entire subset of genes differentially regulated in the absence of *essC* is also similarly altered by loss of *esaB* strongly suggests that EsaB is, like EssC, a core component that is essential for activity of the secretion machinery in strain RN6390.

**Figure 2.**
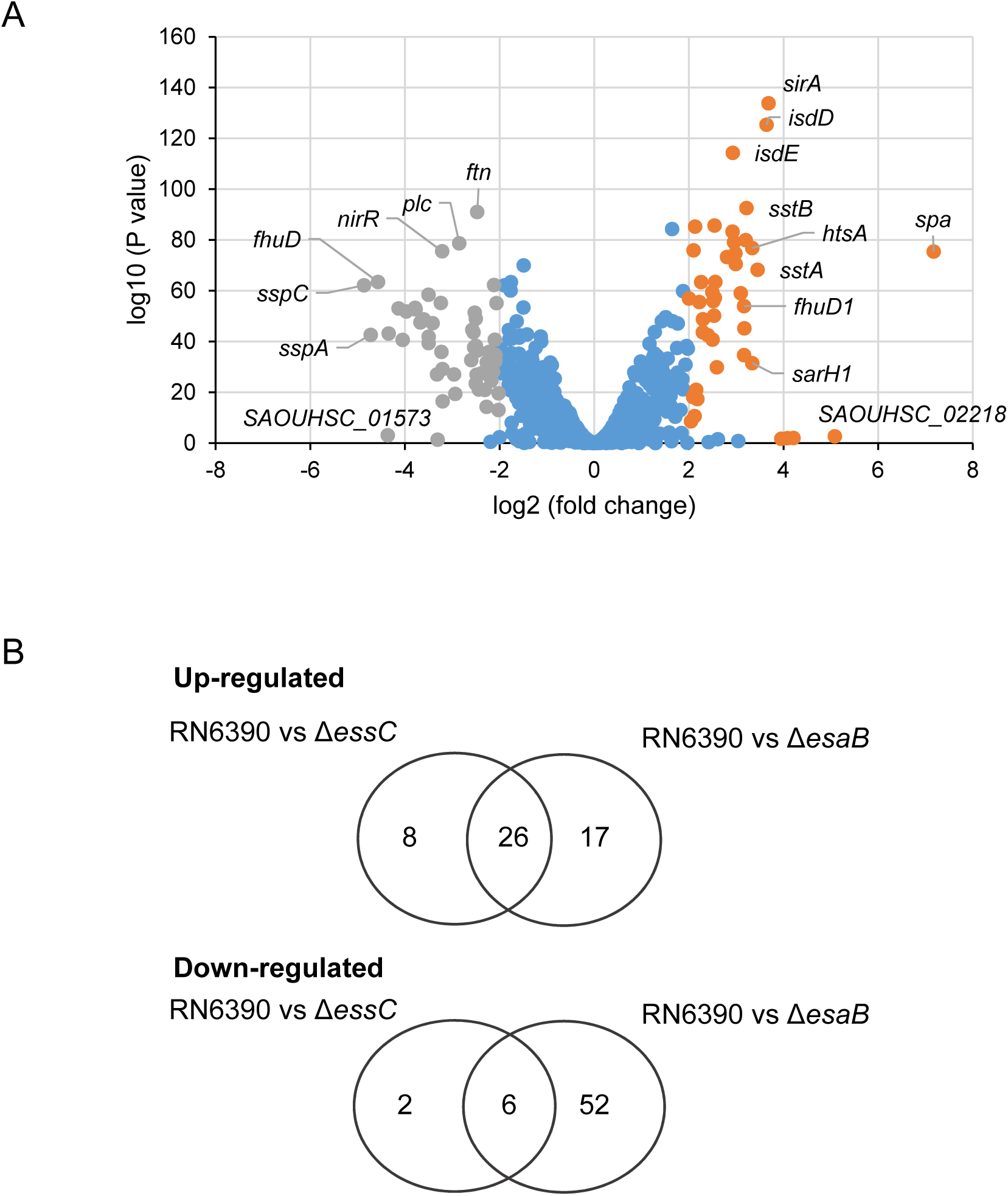
RNA-Seq analysis of differentially regulated genes in the *esaB* mutant strain. (A) Volcano plot representation of the differentially expressed genes in RN6390 strain compared to the isogenic *esaB* mutant. The orange and grey spots represent, respectively, genes that are up- or down-regulated in the *esaB* mutant relative to the parental strain. B. Overlap between up- and down-regulated genes in the *esaB* and *essC* datasets.

**Table 2.** Genes differentially regulated (>log 2 fold) in the RN6390 *esaB* deletion mutant, sorted by ascending fold change (FC). Genes highlighted in grey are also differentially regulated in the *essC* deletion strain. The column on the right shows the fold change (FC) of the same gene in the *essC* dataset where n.s. indicates no statistically significant change in expression level relative to the same gene in the wildtype dataset.

| Locus ID | Gene name | FC in *esaB* mutant | Proposed Function | FC in *essC* mutant |
| --- | --- | --- | --- | --- |
| **Downregulated genes** |  |  |  |  |
| SAOUHSC_00986 | *sspC* | -23.7 | Cysteine protease | n.s. |
| SAOUHSC_00988 | *sspA* | -22.3 | Glutamyl endopeptidase | n.s. |
| SAOUHSC_00987 | *sspB* | -20.8 | Cysteine protease | n.s. |
| SAOUHSC_01573 | — | -19.0 | Unknown, hypothetical protein | n.s. |
| SAOUHSC_01941 | *splB* | -18.8 | Serine protease SplB | -4.3 |
| SAOUHSC_02971 | *aur* | -17.1 | Zinc metalloproteinase aureolysin | n.s. |
| SAOUHSC_01942 | *splA* | -16.4 | Highly specific serine protease specific to *S. aureus* | -5.4 |
| SAOUHSC_02680 | *narH* | -15.7 | Nitrate reductase subunit beta | n.s. |
| SAOUHSC_01944 | — | -14.3 | Unknown, hypothetical protein | -4.5 |
| SAOUHSC_02681 | *narG* | -14.3 | Nitrate reductase subunit alpha | n.s. |
| SAOUHSC_01121 | *hla* | -13.5 | α-hemolysin | -4.1 |
| SAOUHSC_02241 | *lukF* | -13.0 | Unknown, hypothetical protein | -3.3 |
| SAOUHSC_02163 | *hly* | -12.3 | β-hemolysin | n.s. |
| SAOUHSC_01938 | *splD* | -12.2 | Serine protease SplD | -4.3 |
| SAOUHSC_02679 | *narJ* | -12.2 | Nitrate reductase subunit delta | n.s. |
| SAOUHSC_02671 | *narK* | -11.6 | Putative nitrate transporter | n.s. |
| SAOUHSC_02455 | *lacA* | -11.0 | Galactose-6-phosphate isomerase subunit LacA | n.d. |
| SAOUHSC_01530 | — | -10.9 | Hypothetical phage protein | n.s. |
| SAOUHSC_01542 | — | -10.9 | Unknown, SNF2 family protein | n.s. |
| SAOUHSC_01535 | — | -10.9 | Phage capsid protein | n.s. |
| SAOUHSC_02240 | *hly* | -10.5 | Truncated β-hemolysin | n.s. |
| SAOUHSC_02243 | *lukG* | -10.4 | Leukocidin like toxin | -4.5 |
| SAOUHSC_02685 | *nirR* | -10.3 | Unknown, hypothetical protein | n.s. |
| SAOUHSC_01939 | *splC* | -10.3 | Serine protease SplC | -3.2 |
| SAOUHSC_01937 | — | -10.3 | Unknown, hypothetical protein | -2.8 |
| SAOUHSC_02970 | *argR* | -8.8 | Arginine repressor family protein | n.s. |
| SAOUHSC_00113 | *adhE* | -8.6 | Bifunctional acetaldehyde-CoA/alcohol dehydrogenase | n.s. |
| SAOUHSC_00051 | *plc* | -8.1 | 1-phosphatidylinositol phosphodiesterase | -2.5 |
| SAOUHSC_00898 | *argH* | -6.7 | Argininosuccinate lyase | n.s. |
| SAOUHSC_02684 | *nasD* | -6.6 | Assimilatory nitrite reductase [NAD(P)H] large subunit | n.s. |
| SAOUHSC_02709 | *hlyC* | -6.5 | γ-hemolysin component C precursor | -1.8 |
| SAOUHSC_02682 | *nasF* | -6.4 | Uroporphyrin-III C-methyltransferase | n.s. |
| SAOUHSC_02462 | — | -6.4 | Unknown, hypothetical protein | n.s. |
| SAOUHSC_00401 | — | -6.3 | Putative exported protein | -1.6 |
| SAOUHSC_01950 | *epiD* | -6.3 | Flavoprotein | n.s. |
| SAOUHSC_01936 | *splE* | -6.3 | Serine protease SplE | -3.3 |
| SAOUHSC_02454 | *lacB* | -6.3 | Galactose-6-phosphate isomerase subunit LacB | -3.4 |
| SAOUHSC_00899 | *argG* | -6.2 | Argininosuccinate synthase | n.s. |
| SAOUHSC_02108 | *ftn* | -6.1 | Ferritin | n.s. |
| SAOUHSC_00368 | — | -6.1 | Unknown, hypothetical protein | n.s. |
| SAOUHSC_00411 | — | -5.9 | Unknown, hypothetical protein | -2.2 |
| SAOUHSC_01951 | *epiC* | -5.8 | Epidermin biosynthesis protein EpiC | n.s. |
| SAOUHSC_02683 | *nasE* | -5.6 | Assimilatory nitrite reductase [NAD(P)H] small subunit | n.s. |
| SAOUHSC_01935 | *splF* | -5.3 | Serine protease SplF | -2.7 |
| SAOUHSC_02452 | *lacD* | -5.2 | Tagatose 1,6-diphosphate aldolase | -2.6 |
| SAOUHSC_01953 | *epiA* | -5.2 | Gallidermin superfamily EpiA protein | n.s. |
| SAOUHSC_02941 | *nrdG* | -4.9 | Anaerobic ribonucleotide reductase activating protein | n.s. |
| SAOUHSC_00717 | *saeP* | -4.7 | Putative lipoprotein | -1.4 |
| SAOUHSC_01990 | *glnQ* | -4.6 | Glutamine transport ATP-binding protein | n.s. |
| SAOUHSC_02557 | *yut* | -4.5 | Putative urea transporter | n.s. |
| SAOUHSC_01949 | *epiP* | -4.4 | Intracellular serine protease | n.s. |
| SAOUHSC_00120 | *capG* | -4.4 | UDP-N-acetylglucosamine 2-epimerase | n.s. |
| SAOUHSC_01952 | *bsaB* | -4.4 | Lantibiotic epidermin biosynthesis protein EpiB | n.s. |
| SAOUHSC_03015 | *hisZ* | -4.4 | ATP phosphoribosyltransferase regulatory subunit | n.s. |
| SAOUHSC_00119 | *capF* | -4.4 | Capsular polysaccharide biosynthesis protein CapF | n.s. |
| SAOUHSC_02463 | *hysA* | -4.3 | Hyaluronate lyase | n.s. |
| SAOUHSC_02453 | *lacC* | -4.1 | Tagatose-6-phosphate kinase | -2.2 |
| SAOUHSC_00608 | *adh1* | -4.1 | Alcohol dehydrogenase | n.s. |
| **Upregulated genes** |  |  |  |  |
| SAOUHSC_02767 | *opp-1A* | 4.0 | Peptide ABC transporter substrate-binding protein | 2.6 |
| SAOUHSC_02655 | — | 4.2 | Unknown, hypothetical protein | 6.3 |
| SAOUHSC_01292 | — | 4.4 | Putative DNA-binding protein | n.s. |
| SAOUHSC_00130 | *isdI* | 4.4 | Heme-degrading monooxygenase IsdI | 5.7 |
| SAOUHSC_00176 | — | 4.5 | Extracellular solute-binding protein | n.s. |
| SAOUHSC_02435 | *sfaA* | 4.5 | Putative transporter | 6.7 |
| SAOUHSC_02799 | *sarT* | 4.6 | Staphylococcal accessory regulator | n.s. |
| SAOUHSC_02432 | — | 4.8 | Unknown, hypothetical protein | 6.2 |
| SAOUHSC_02245 | — | 4.9 | Unknown, hypothetical protein | 6.5 |
| SAOUHSC_00652 | *fhuA* | 5.1 | Ferrichrome ABC transporter ATP-binding protein FhuA | 7.0 |
| SAOUHSC_00071 | *sirC* | 5.3 | Involved in staphyloferrin B transport into the cytoplasm | 4.6 |
| SAOUHSC_00131 | — | 5.3 | Putative membrane protein | 6.1 |
| SAOUHSC_02821 | — | 5.8 | Putative membrane protein | n.s. |
| SAOUHSC_02719 | — | 6.2 | ABC transporter ATP-binding protein | 5.5 |
| SAOUHSC_01920 | — | 6.3 | Putative lipoprotein | n.s. |
| SAOUHSC_02428 | *htsB* | 6.3 | Heme transport system permease HtsB | 5.4 |
| SAOUHSC_00974 | — | 6.4 | Unknown, hypothetical protein | n.s. |
| SAOUHSC_01081 | *isdA* | 6.5 | Iron-regulated heme-iron binding protein | 5.4 |
| SAOUHSC_00072 | *sirB* | 6.5 | Involved in staphyloferrin B transport into the cytoplasm | 7.4 |
| SAOUHSC_02554 | *fhuD2* | 6.5 | Ferric hydroxamate receptor 1 FhuD2 | 6.8 |
| SAOUHSC_01090 | — | 6.7 | Unknown, hypothetical protein | 3.9 |
| SAOUHSC_00973 | — | 7.9 | Unknown, hypothetical protein | n.s. |
| SAOUHSC_01086 | *isdF* | 8.5 | ABC permease IsdF | 6.1 |
| SAOUHSC_01085 | *isdE* | 8.6 | Heme-receptor lipoprotein IsdE | 5.6 |
| SAOUHSC_01089 | *isdG* | 8.7 | Heme-degrading monooxygenase IsdG | 4.7 |
| SAOUHSC_01087 | — | 8.9 | Iron compound ABC transporter permease | 6.3 |
| SAOUHSC_01082 | *isdC* | 8.9 | Heme transporter IsdC | 5.5 |
| SAOUHSC_00748 | *sstC* | 9.6 | Ferrichrome ABC transporter ATP-binding protein SstC | 9.1 |
| SAOUHSC_00545 | *sdrD* | 10.0 | Fibrinogen-binding protein SdrD | n.s. |
| SAOUHSC_02246 | *fhuD1* | 10.0 | Iron compound ABC transporter FhuD1 | 8.0 |
| SAOUHSC_00972 | — | 10.1 | Unknown, hypothetical protein | n.s. |
| SAOUHSC_01088 | *srtB* | 10.2 | Sortase StrB | 6.2 |
| SAOUHSC_00747 | *sstB* | 10.4 | Ferrichrome ABC transporter permease SstB | 9.0 |
| SAOUHSC_00070 | *sarH1* | 11.2 | Unknown, hypothetical regulatory-like protein | n.s. |
| SAOUHSC_02430 | *htsA* | 11.2 | Heme transport system lipoprotein HtsA | 10.5 |
| SAOUHSC_00746 | *sstA* | 11.9 | Ferrichrome ABC transporter permease SstA | 10.9 |
| SAOUHSC_01084 | *isdD* | 13.3 | ATP-hydrolysing and heme-binding protein IsdD | 6.2 |
| SAOUHSC_00074 | *sirA* | 13.6 | Receptor component of staphyloferrin B | 16.3 |
| SAOUHSC_01514 | — | 15.6 | Unknown, hypothetical protein | n.s. |
| SAOUHSC_02232 | — | 16.7 | Hypothetical phage protein | n.s. |
| SAOUHSC_02084 | — | 17.7 | Phage repressor protein | n.s. |
| SAOUHSC_02218 | — | 25.9 | Unknown, hypothetical protein | n.s. |
| SAOUHSC_00069 | *spa* | 51.5 | Protein A | n.s. |

As mentioned above, a subset of transcripts were differentially expressed in the *esaB* but not the *essC* strain. These include downregulated genes required for anaerobic nitrate respiration (*narGHJ*/*narK*), some secreted proteases (*sspA/B/C*, *aur*), capsular polysaccharide synthesis (*capG/F/hysA*), lactose metabolism (*lacB/C/D*) and antimicrobial peptide synthesis (*epiA/C/D/P*). Many of these genes are under control of the essential two component regulatory system AirSR (formerly YhcSR) (25-28). These findings suggest that EsaB may have additional roles in the cell in addition to its requirement for T7 protein secretion.

### EsaB is present at low amounts in cells of *S. aureus* RN6390

To explore the biological role of EsaB in T7 secretion, we firstly overproduced recombinant EsaB with a cleavable His-tag in *E. coli*, and following cleavage of the tag the protein was further purified by gel filtration chromatography (Fig 3A, B). The purified protein, which eluted with an estimated molecular mass of approximately 12.8 kDa, is close to the expected size of a monomer (9.1 kDa + 0.3 kDa retained following cleavage of the tag = 9.4 kDa). This is in agreement with structural analysis of the *B. subtilis* EsaB homologue, YukD, which also appears to be monomeric (20).

**Figure 3.**
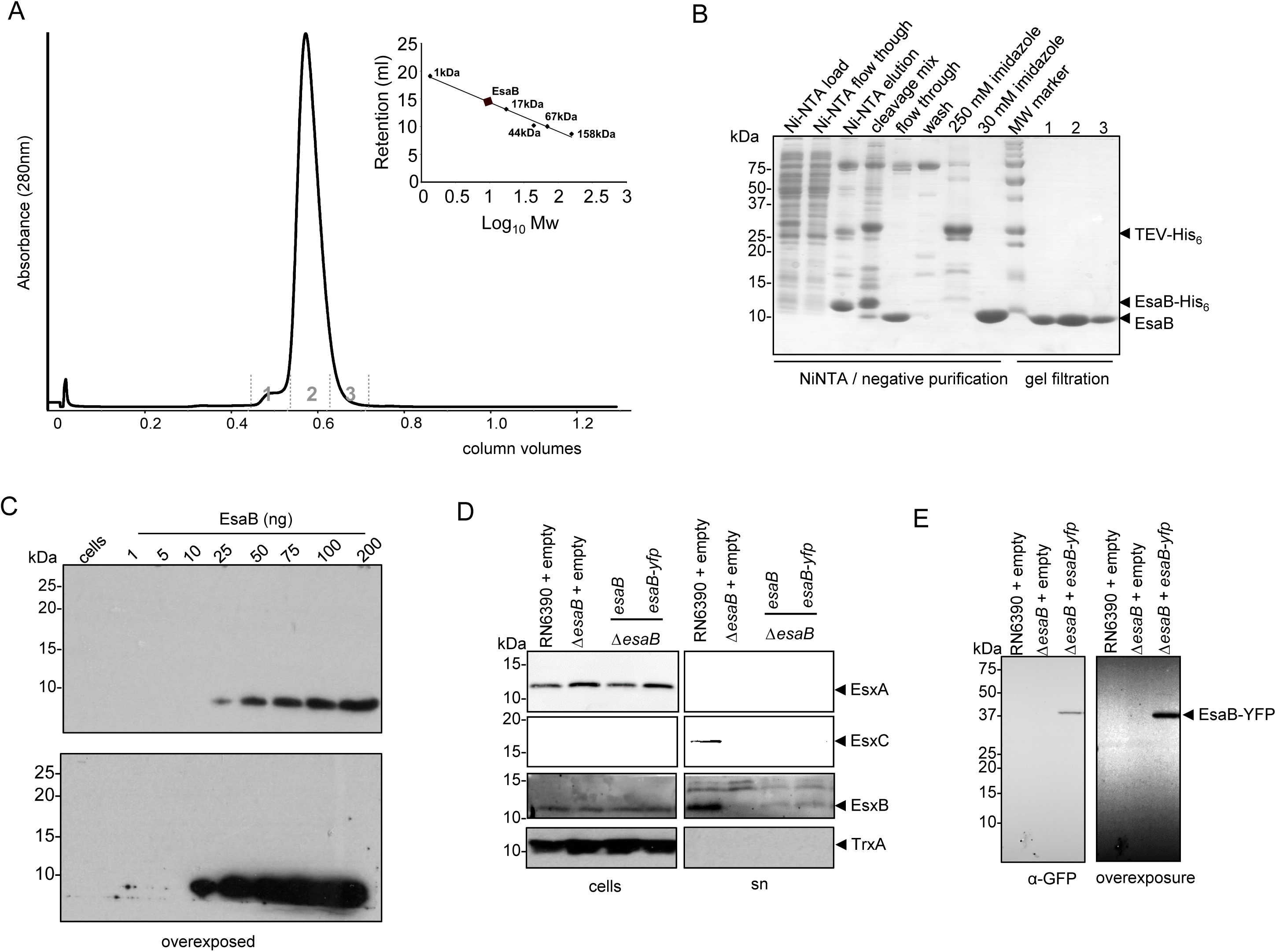
EsaB is present in cells at low amounts. (A) Purification of EsaB by gel filtration chromatography. Inset shows column calibration. (B) SDS PAGE analysis of EsaB during purification steps. Lanes labelled 1, 2 and 3 correspond to similarly-labelled fractions from the gel filtration column. (C) Titration of α-EsaB antibodies. The indicated amounts of purified EsaB, alongside 30 μl of OD_600_ 5 adjusted cells were loaded on a SDS-PAGE as indicated and blotted using α-EsaB antibodies. Two exposures of the blot are shown. (D) RN6390 harbouring empty pRAB11, and the isogenic *esaB* deletion strain harbouring pRAB11, or pRAB11 encoding native EsaB or EsaB-YFP was cultured aerobically in TSB medium until an OD_600_ of 2 was reached. Samples were fractionated to give cells and supernatant (sn), and supernatant proteins were precipitated using TCA. For each gel, 10 μl of OD_600_ 1 adjusted cells and 15 μl of culture supernatant were loaded. Blots were probed with anti-EsxA, anti-EsxB or anti-EsxC antisera, alongside anti-TrxA (cytoplasmic control). Cell and supernatant samples have been blotted on the same gel but intervening lanes have been spliced out. (E) EsaB-YFP can be detected in whole cells. RN6390 harbouring empty pRAB11, and the isogenic *esaB* deletion strain harbouring pRAB11, or pRAB11 encoding EsaB-YFP was cultured aerobically in TSB medium until an OD_600_ of 2 was reached. Whole cell samples (20 μl of OD_600_ 2 adjusted cells) were loaded and blots were probed with anti GFP antibodies. Two exposures of the blot are shown.

Polyclonal antisera were raised against purified EsaB and the antibody was affinity purified against the EsaB antigen, before being used to detect the protein in whole cells of *S. aureus*. Fig 3C shows that although the purified antiserum could clearly recognize purified EsaB, it did not detected a band of the expected size of EsaB in whole cells. Probing a dilution series of purified EsaB indicated that the antibody was able to cross-react with as little as 25ng of protein, which is equivalent to 1.6 x 10^11^ EsaB molecules. Since the antibody was unable to detect EsaB in whole cells from 9.6 x 10^8^ colony forming units that were loaded onto the SDS gel, we conclude that are less than 170 molecules of EsaB per cell.

Since we were unable to detect native EsaB in *S. aureus* cell extracts, we constructed a series of tagged variants for which commercial antisera were available. To this end we introduced His_6_, Myc, hemagglutinin (HA) and Strep epitopes onto the N-terminus of EsaB, and His_6_, Myc, HA, mCherry or FLAG epitopes onto the C-terminus, but in each case were unable to detect the tagged protein (not shown). We also introduced His_6_ and His_9_ epitopes into two predicted loop regions internal to the EsaB sequence but again were unable to detect tagged EsaB (not shown). The only tag we introduced that allowed detection of EsaB was a C-terminal yellow fluorescent protein (YFP) tag. Fig 3D shows that basal production of either native (untagged) EsaB or EsaB-YFP from plasmid vector pRAB11 was sufficient to restore secretion of the T7SS extracellular protein EsxA and of substrates EsxB and EsxC to the culture supernatant. Blotting the same cell samples for the presence of the YFP fusion protein (Fig 3E) showed that it migrated at close to the predicted mass (37 kDa) of the EsaB fusion. There was no evidence for degradation of the fusion protein even after prolonged exposure of the immunoblot (Fig 3E). We conclude that the YFP-tagged variant of EsaB probably retains functionality.

### EsaB-YFP partially localizes to the cell membrane

EsaB is predicted to be a soluble cytoplasmic protein (10), and is known to share structural homology with ubiquitin (20). Interestingly, a domain sharing the same fold is also associated with the actinobacterial T7SS, being found at the cytoplasmic N-terminus of EccD (29), indicating that ubiquitin-like proteins are essential features of all T7SSs. To determine the subcellular location of EsaB-YFP, we blotted secreted and whole cell samples of the *esaB* mutant strain producing plasmid-encoded EsaB-YFP with the YFP antiserum. Fig 4A shows that EsaB-YFP was associated exclusively with the cellular fraction.

**Figure 4.**
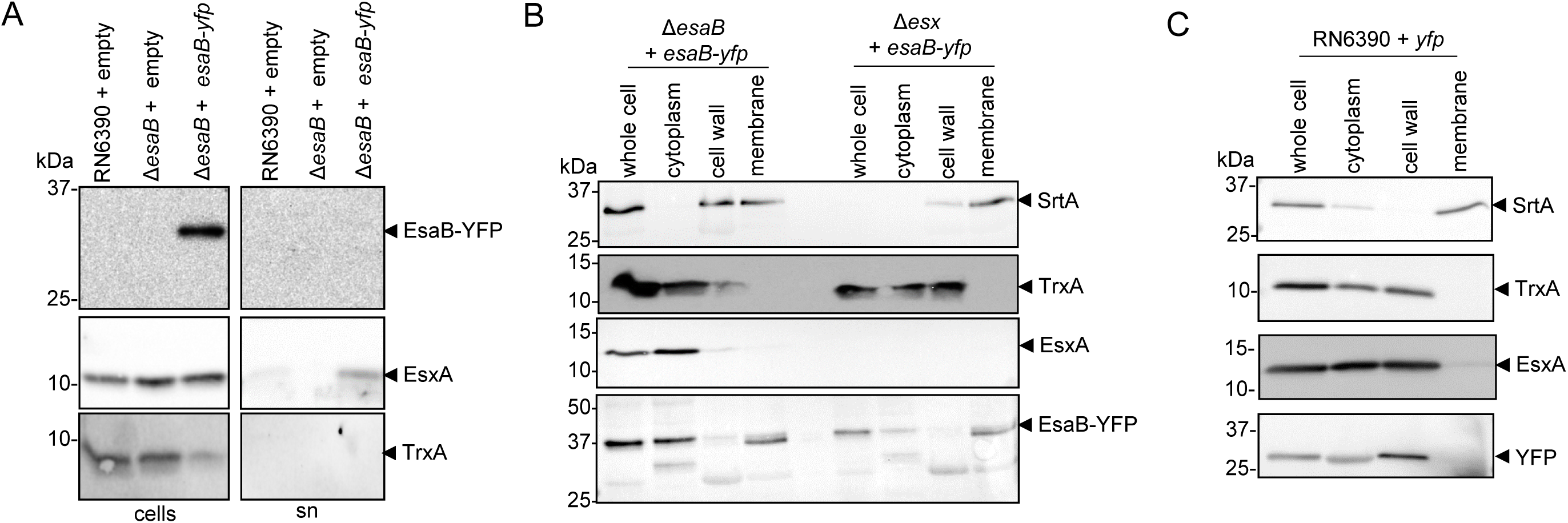
EsaB-YFP localizes to the cytoplasm and membrane. (A) EsaB-YFP is not secreted in *S. aureus* strain RN6390. RN6390 harbouring empty pRAB11 and the isogenic *ΔesaB* strain harbouring empty pRAB11 or pRAB11 encoding EsaB-YFP were cultured in TSB medium until mid-log phase and separated into cellular and supernatant fractions (sn). For each gel, 10 μl of OD_600_ 1 adjusted cells and 15 μl of TCA-precipitated culture supernatant were loaded. Blots were probed with anti-EsxA, anti-TrxA (cytoplasmic control) and anti-GFP antisera. Cell and supernatant samples have been blotted on the same gel but intervening lanes have been spliced out. Subcellular localisation of (B) EsaB-YFP in RN6390 and an isogenic Δ*esx* (Δ(*esxA*-*esaG*) strain or (C) YFP in RN6390. Cells were grown aerobically in TSB to mid-log phase and fractionated as indicated in the Methods. Equivalents amount of each fraction was probed with anti-TrxA (cytoplasmic control), anti-SrtA (membrane control), anti-EsxA and anti-GFP antisera.

We next fractionated these cells to obtain cytoplasm, cell wall and membrane fractions. Immunoblotting with antisera to control proteins known to localize to the cell membrane (SrtA) and cytoplasm (TrxA) indicated that the fractionation had been largely successful, although some SrtA was found in the cell wall fraction (Fig 4B). Blotting these same fractions for the presence of EsaB-YFP showed that the protein localised to both the cytoplasm and membrane fractions. Some degradation of the fusion protein was also noted in these experiments which may result from the activation of proteases during fractionation. When unfused YFP was produced in the wild type strain it did not localise to the membrane (Fig 4C), indicating that membrane binding was unlikely to be mediated through the YFP portion of the fusion.

Next we tested whether EsaB-YFP localised to the membrane through interactions with membrane components of the T7SS. To this end we repeated the fractionation in a strain carrying a chromosomal deletion in all twelve genes at the *ess* locus (Fig 1A). However this did not alter the localization of EsaB-YFP, which was still detected in both cytoplasm and membrane fractions (Fig 4B). It may be that EsaB-YFP localises to the membrane through interaction with additional membrane proteins, consistent with additional, T7SS-independent roles for EsaB suggested through the RNA-Seq analysis. Alternatively, we cannot rule out that the membrane localization arises as an artifact of the C-terminal YFP tag, since this tag is known to influence protein behaviour (e.g. (30)).

### Mutagenesis of conserved hydrophilic and hydrophobic patches on EsaB

An alignment of EsaB homologues encoded across firmicutes (Fig 5A) identifies a number of highly conserved amino acids. Many of these are hydrophilic and fall on one face of the predicted structure of EsaB including T8 (*S. aureus* numbering) which is highly conserved as either threonine or serine, and the invariant K56. The presence of an invariant lysine is particularly intriguing since there are a number of highly conserved lysine residues on the structurally-related protein ubiquitin, that are used to assemble polyubiquitin chains (31). To probe potential roles of these conserved residues we mutated each of T8, D10, L21, K30, K52, K56, L66, G74 and D75 to alanine on plasmid-encoded EsaB and assessed whether the variant EsaB proteins were able to restore T7 secretion activity to the *esaB* deletion strain.

**Figure 5.**
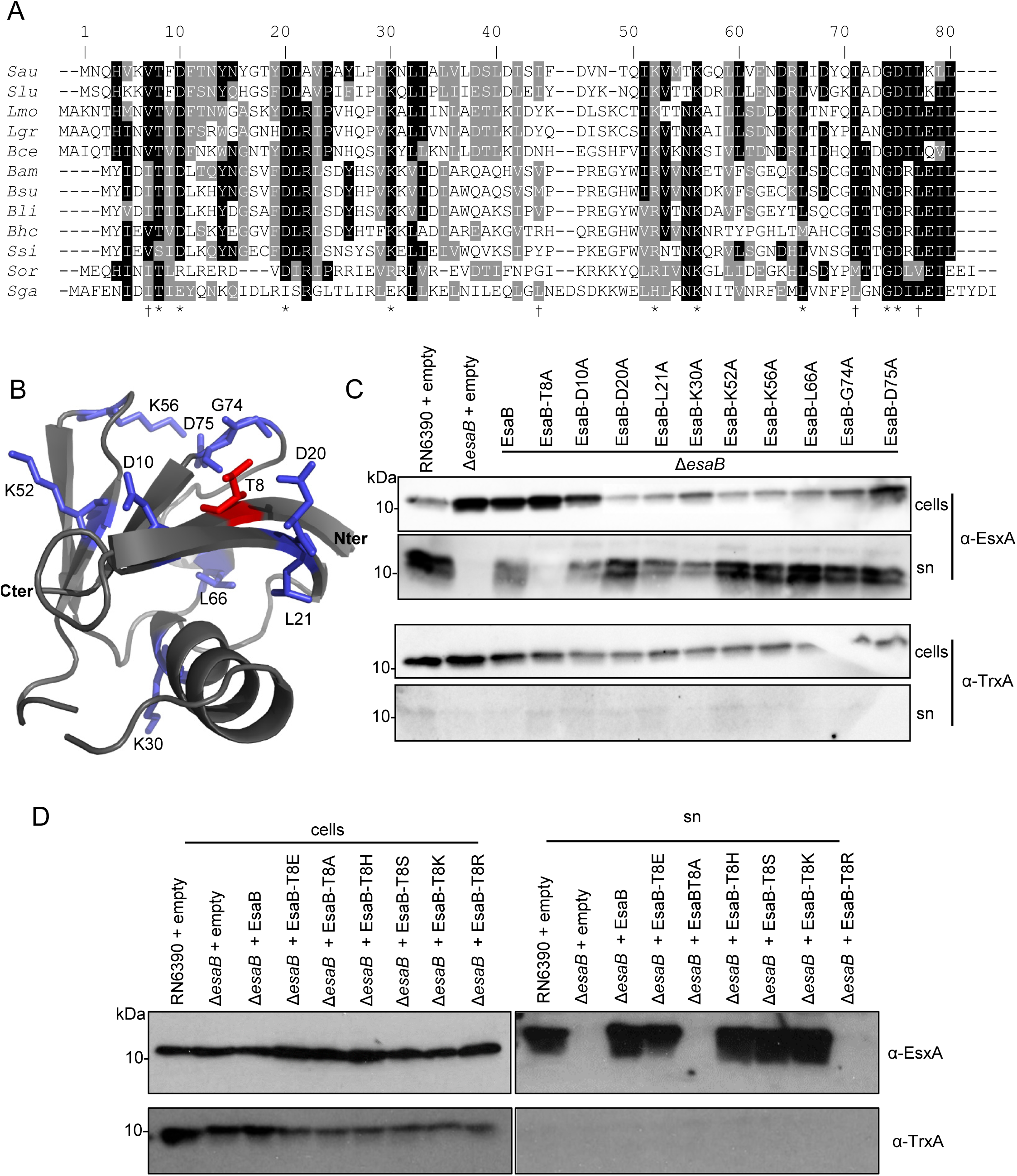
Site-directed mutagenesis of conserved residues of EsaB. (A) Sequence alignment of EsaB homologues from: Sau - *Staphylococcus aureus*; Slu - *Staphylococcus lugdunensis*; Lmo - *Listeria monocytogenes*; Lgr - *Listeria grayi*; Bce - *Bacillus cereus*; Bam - *Bacillus amyloliquefaciens*; Bsu - *Bacillus subtilis*; Bli - *Bacillus licheniformis*; Bhc - *Bhargavaea cecembensis*; Ssi - *Solibacillus silvestris*; Sor - *Streptococcus oralis*; Sga - *Streptococcus gallolyticus*. * indicate conserved residues and † indicates residues forming a potential hydrophobic patch that were mutated in this work. (B) Model of *S. aureus* EsaB with positions of conserved residues targeted for mutagenesis highlighted. The N- and C-termini are also indicated. (C) and (D) RN6390 harbouring empty pRAB11, and the isogenic *esaB* deletion strain harbouring pRAB11, or pRAB11 encoding native the indicated variants of EsaB were cultured aerobically in TSB medium until an OD_600_ of 2 was reached. Samples were fractionated to give cells and supernatant (sn), and supernatant proteins were precipitated using TCA. For each gel, 10 μl of OD_600_ 1 adjusted cells and 15 μl of culture supernatant were loaded. Blots were probed with anti-EsxA, and anti-TrxA (cytoplasmic control) antisera.

Fig 5C shows that alanine substitutions of each of these conserved residues was tolerated by EsaB with the exception of T8A, which completely abolished EsaB activity. To test whether other side chain substitutions were permissive at T8, we subsequently constructed EsaB T8S, T8E, T8H, T8K and T8R substitutions. As seen in Fig 5D, in addition to T8A the T8R substitution also abolished EsaB activity, but the other substitutions resulted in active protein.

Ubiquitin has a conserved hydrophobic patch (Fig 6A, *left*) that forms a common site of interaction with many different binding partners (32). Analysis of the predicted structure of EsaB (Fig 6A, *right*) shows that there are some hydrophobic residues on the surface potentially at positions approximating the hydrophobic patch region of ubiquitin. To assess whether these hydrophobic residues may be involved in EsaB function, we firstly mutated V7, I44, I71 and L77 to alanine residues. Fig 6B shows that these substitutions did not detectably affect T7 secretion indicating that the function of EsaB had not been compromised. We next substituted each of these residues for a positively-charged lysine. This more drastic change of amino acid side-chain was still tolerated at positions 44 and 77, but inactivated EsaB when substituted for V7 or I71.

**Figure 6.**
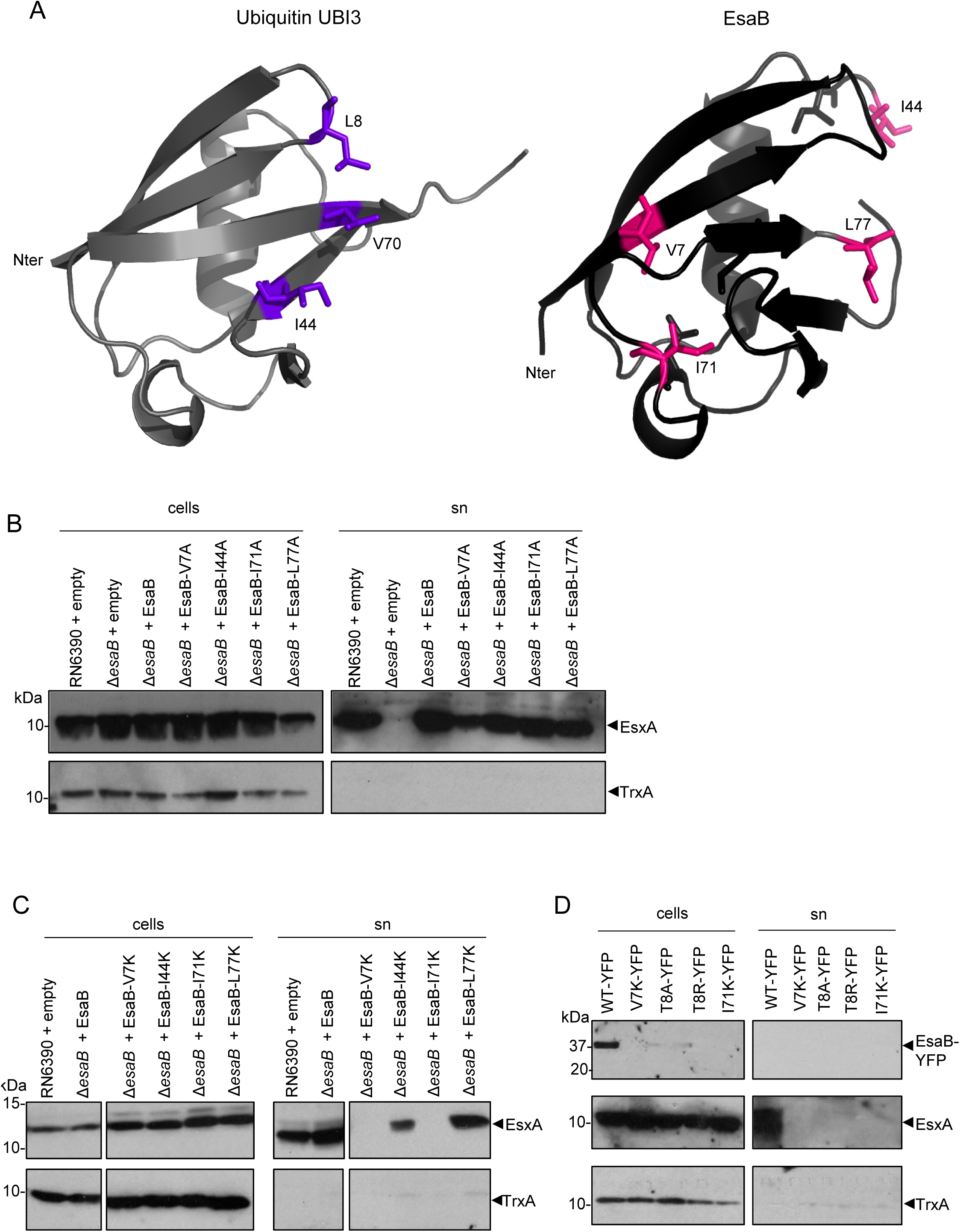
Mutagenesis of a hydrophobic patch on EsaB. (A) Ribbon model of ubiquitin (left; PDB: 1ubi) with residues forming a conserved hydrophobic patch highlighted in purple and *S. aureus* EsaB (right) with positions of hydrophobic residues targeted for mutagenesis shown in pink. (B) and (C) RN6390 harbouring empty pRAB11, and the isogenic *esaB* deletion strain harbouring pRAB11, or pRAB11 encoding native EsaB or the indicated amino acid-substituted variants were cultured aerobically in TSB medium until an OD_600_ of 2 was reached. Samples were fractionated to give cells and supernatant (sn), and supernatant proteins were precipitated using TCA. For each sample, 10 μl of OD_600_ 1 adjusted cells and 15 μl of culture supernatant were loaded. Blots were probed with anti-EsxA, and anti-TrxA (cytoplasmic control) antisera. Cell and supernatant samples have been blotted on the same gel but intervening lanes have been spliced out. (D) The Δ*esaB* strain harbouring pRAB11 encoding EsaB-YFP (WT-YFP) or the indicated amino acid-substituted variants were cultured and fractionated as in (B) and (C). For each sample, 10 μl of OD_600_ 1 adjusted cells and 15 μl of culture supernatant were loaded and blots were probed with anti-EsxA, anti-TrxA or anti-GFP antisera.

Finally we attempted to assess whether any of the inactivating substitutions E7K, T8A, T8R or I71K altered the subcellular location of EsaB-YFP. However, when we introduced each of these substitutions into EsaB-YFP we found that they destabilised the protein as it was almost undetectable in whole cells (Fig 6D), precluding further analysis. We are therefore unable to determine whether these substitutions directly alter EsaB function or have an indirect effect by disrupting folding.

## DISCUSSION

In this work we have investigated the role of EsaB in Type VII secretion. EsaB proteins are conserved in firmicutes that produce the T7SS and are encoded at the same loci. Previous work had implicated EsaB in the regulation of *esxC* transcripts (11), although this cannot be a conserved role for EsaB proteins as they are found in all *S. aureus* strains, including the subset that do not encode *esxC* (16). Here we show that EsaB does not regulate *esxC* in strain RN6390, nor any of the other genes encoded at the *ess* locus. Instead, deletion of *esaB* is associated with upregulation of genes involved in iron acquisition, mirroring the upregulation of iron-acquisition genes seen when the core T7 component, EssC, is absent (15). This supports the notion that EsaB is a core component of the secretion machinery in RN6390, and in agreement with this, deletion of *esaB* prevented export of the T7-dependent extracellular proteins EsxA, EsxB and EsxC. This conclusion is also in agreement with related studies in *B. subtilis*, where the EsaB homologue YukD was shown to be essential for secretion of the WXG100 protein YukE (17, 18).

The precise role of EsaB in T7 secretion is unclear. Structural analysis of *B. subtilis* YukD shows that it shares a very similar fold to ubiquitin but that it lacks the ability to be conjugated with other proteins (20). Interestingly, a ubiquitin-like domain is also associated with the actinobacterial T7SS, being found at the cytoplasmic N-terminus of the polytopic EccD membrane component (29), suggesting that EsaB-like components are essential features of all T7SSs. Ubiquitin interacts directly with many different protein binding partners (32), and it is therefore likely that EsaB interacts with one or more components of the T7SS, potentially regulating activity. Post-translational regulation of the *S. aureus* T7SS has been suggested because in some growth conditions the secretion machinery is present but there is no or very little substrate secretion (12, 19). Other protein secretion systems are also post-translationally regulated, for example the flagellar type III secretion system is regulated through interaction of the FliI component with the second messenger cyclic di-GMP (33), and Type VI secretion systems are regulated by phosphorylation (34). In this context, EsaB proteins contain a highly conserved threonine (or serine) residue close to their N-termini which we considered as a potential site for phosphorylation. Intriguingly, substitution of EsaB T8 for alanine abolished the function of EsaB, although introduction of either the phospho-mimetic glutamate at this position or a positively charged lysine did not affect EsaB activity.

The low cellular levels of EsaB precluded further analysis of the native protein, but a C-terminal fusion of EsaB with YFP partially localised to the cell membrane. We reasoned that binding of EsaB-YFP to membranes was mediated through interaction with one or more of the T7SS membrane proteins. However, some EsaB-YFP was still membrane associated when it was analysed in a strain lacking all of the core T7 components, suggesting that it may interact with additional membrane proteins potentially unrelated to the T7SS. Support for this suggestion comes from RNA-Seq analysis of the *esaB* mutant strain. In addition to a common set of genes showing similar regulation in the *esaB* and *essC* strains, a further subset of genes were uniquely deregulated in the *esaB* mutant. Many of the genes in this EsaB-specific subset are part of the AirSR regulon (25-28). The AirSR two component system responds to oxidation signals via a redox-active [2Fe-2S] cluster in the sensor kinase AirS to regulate diverse sets of genes involved in anaerobic respiration, lactose metabolism and capsule biosynthesis. In future it will be interesting to further decipher the roles of EsaB in T7 protein secretion and *S. aureus* physiology.

## ACKNOWLEDGEMENTS

This study was supported by the Wellcome Trust (through Investigator Award 10183/Z/15/Z to TP and through Clinical PhD studentship support to CPH through grant 104241/z/14/z), the Biotechnology and Biological Sciences Research Council and the Medical Research Council (through grants BB/H007571/1 and MR/M011224/1, respectively). We thank Dr Sarah Murdoch for calculating the relationship between OD 600nm and CFU for RN6390. Dr Francesca Short is thanked for her assistance with RNA-Seq data analysis and Professor Nicola R. Stanley-Wall for her advice with RNA extraction. The authors declare no conflicts of interest.

---

**Table S1.**
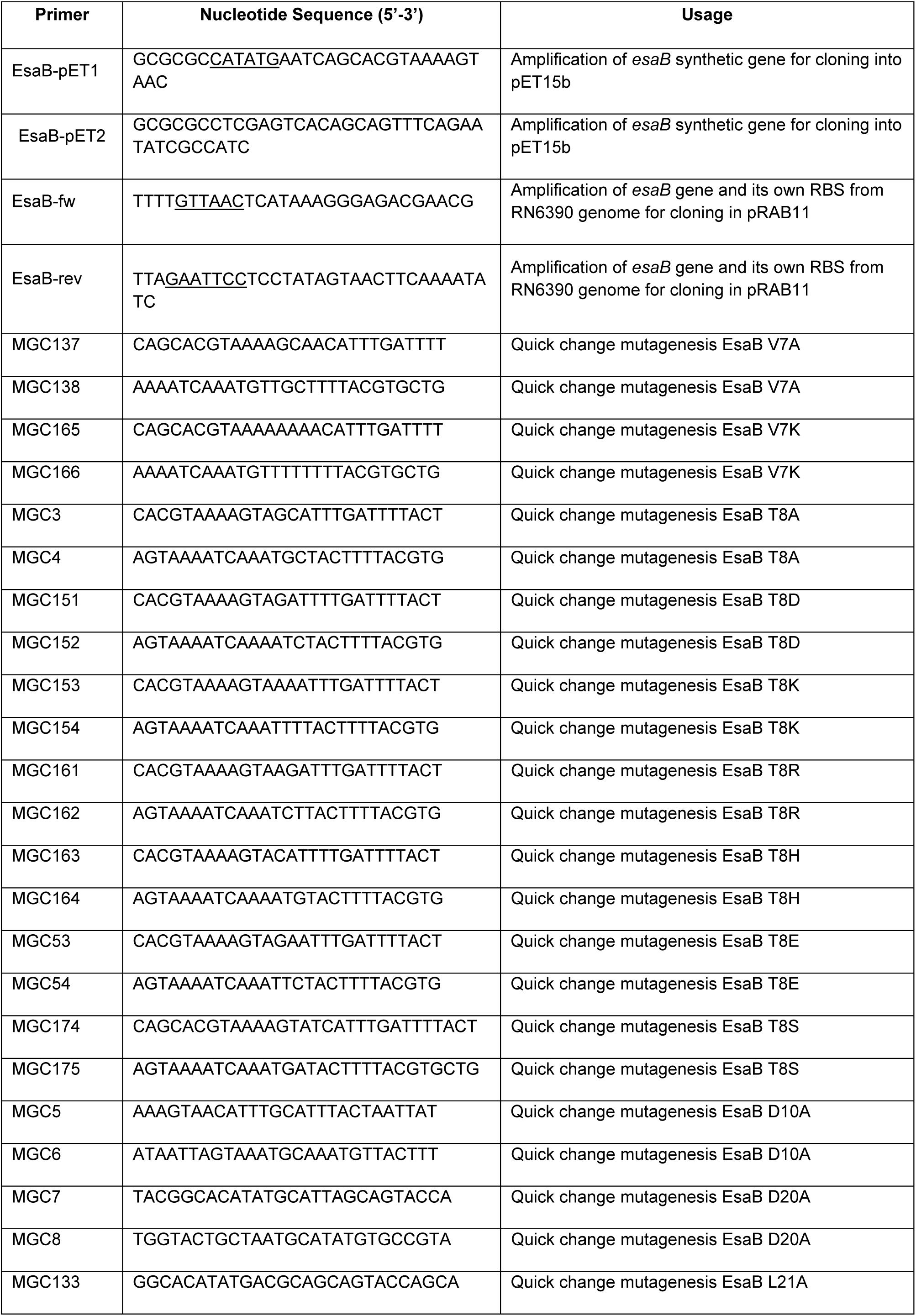
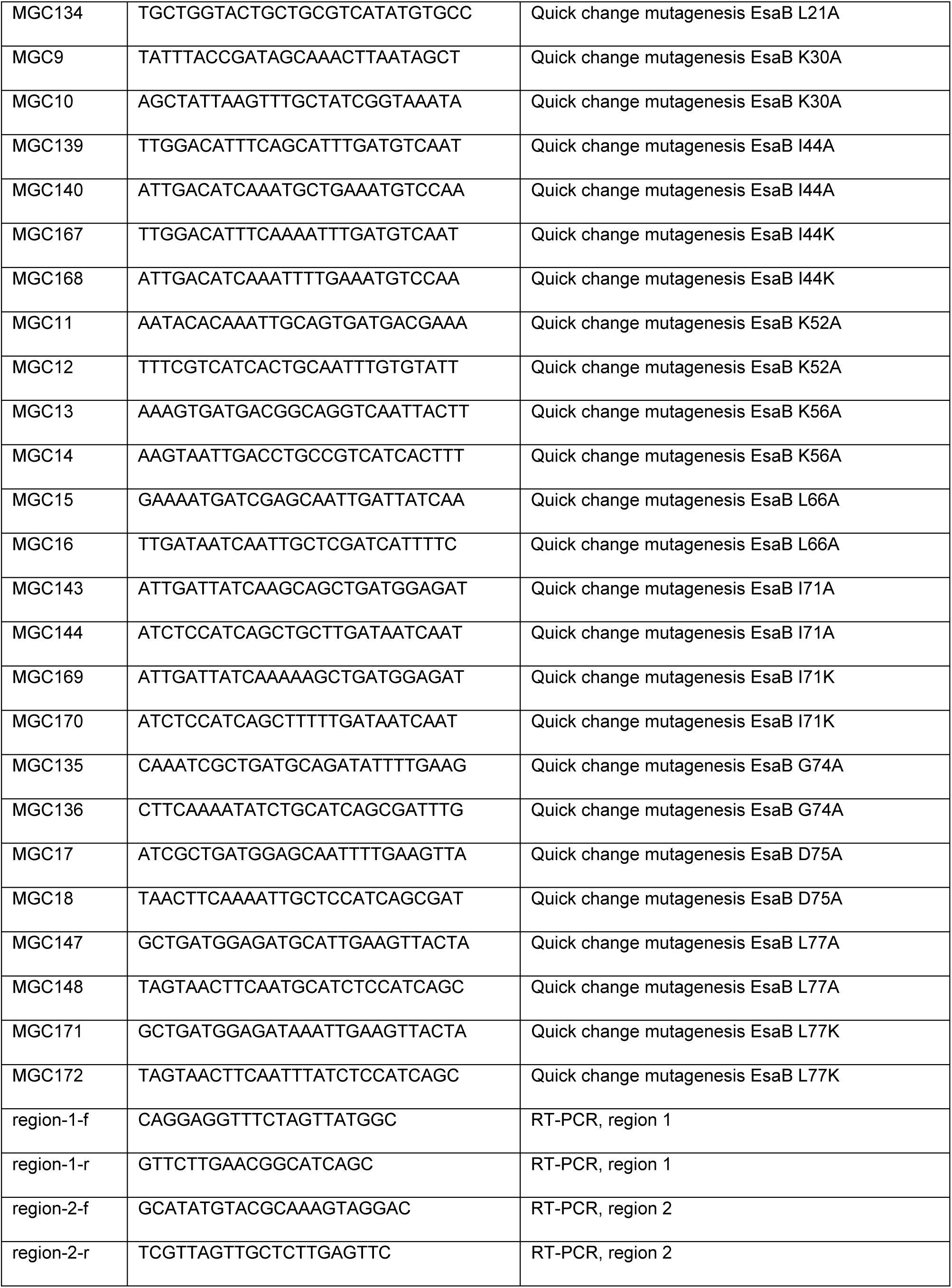
Oligonucleotides used in this study. Restriction enzyme sites have been underlined.

